# A New Look at Medaka: Surprising Deep Diving Behavior Uncovered in the Lab

**DOI:** 10.1101/2025.08.12.669509

**Authors:** Shoji Fukamachi, Tamaki Uchikawa, Eiji Watanabe

## Abstract

Medaka (*Oryzias latipes* and *O. sakaizumii*) are small freshwater fish native to East Asia, commonly found in shallow waters. However, their presence and behavior in deeper habitats remain poorly understood due to observational challenges. Here, we introduce a novel experimental system—a 9-meter-tall cylindrical tank made from vertically connected transparent PVC pipes—that enables direct observation of medaka diving behavior. Our results demonstrate that medaka voluntarily dive to depths of up to 9 meters and remain at the bottom for extended periods, a behavior contrasting with that of zebrafish (*Danio rerio*), which rarely reached such depths. Moreover, when fish were introduced from the bottom of the tank, zebrafish exhibited repeated vertical movements, ascending and descending but rarely reaching the water surface, whereas medaka consistently remained at the bottom. While primarily descriptive, these findings represent a significant and previously unrecognized aspect of medaka ecology, challenging the traditional view of them as purely surface-dwelling fish. This study provides a unique tool and foundational insight that opens new avenues for ecological and physiological investigations of fish depth preferences.

## Introduction

Medaka (*Oryzias latipes* and *O. sakaizumii*) are small teleost fishes native to Japan, Korea, and China. In Japan, they are commonly observed in shoals from spring to autumn in shallow waters such as slow-flowing rivers and rice paddies. In winter, however, they vanish from these shallow habitats, implying a seasonal migration to deeper waters for overwintering. Capture surveys conducted during winter in agricultural waterways support this view, showing a positive relationship between water depth and the number of medaka caught (Ishikawa & Azuma, 2005; Minagawa et al., 2010). Yet, the maximum depths of these artificial channels are only about 30–60 cm—likely much shallower than the deepest habitats medaka can occupy in nature. It therefore remains unknown whether medaka are capable of, and perhaps prefer, overwintering in habitats several meters deep.

Medaka, with their upward-facing mouths, are generally considered surface-oriented swimmers and foragers, and indeed are frequently observed in shallow surface waters in natural habitats. However, systematic observations from offshore environments or from deeper strata (midwater or near-bottom layers) are lacking, leaving their vertical distribution in these zones largely undocumented. Nevertheless, in aquarium tanks approximately 30 cm deep, medaka have been observed not only just beneath the surface—unlike obligate surface dwellers such as hatchetfish—but also in midwater and near-bottom zones while actively foraging, suggesting that their depth use may be more flexible than commonly assumed.

In the course of conducting an ecological survey—supported in part by artificial intelligence tools—of wild medaka inhabiting the Nakaikemi Wetland (Fukui prefecture, Japan)—where the maximum water depth reaches approximately 4 meters (Murakami et al., 2021)—we faced a practical need to determine the depth range that should be included in our investigations. Although previous studies using artificially elevated water pressure have demonstrated that medaka can survive and behave normally at pressures equivalent to 300 meters depth (3 MPa) (Iwahashi et al., 2007), it is unlikely that such extreme depths represent their natural habitat. Therefore, to assess how deep medaka will voluntarily dive—and specifically whether they can inhabit depths of up to 4 meters—we conducted experiments using a tall, cylindrical tank composed of vertically connected PVC pipes. While the findings presented here are relatively simple and primarily descriptive, they offer novel insights into the depth ecology of this model organism, filling a notable gap in its natural history and potentially informing both ecological research and conservation efforts.

## Materials and Methods

### Fish used

For *Oryzias latipes*, we used the d-rR strain, a commonly employed laboratory strain. For *O. sakaizumii*, we used wild individuals collected from the Nakaikemi Wetland (hereafter referred to as “Nakaikemi”). All medaka were reared from fertilized eggs to adulthood in the laboratory at approximately 25 °C under a 14 h light:10 h dark photoperiod. Prior to the experiments, individuals were kept outdoors for several weeks to several months. For zebrafish (*Danio rerio*), we used commercially obtained adults that were acclimated in the laboratory for two days before the experiments. All fish used were sexually mature at the time of testing.

### Apparatus and procedure

Transparent PVC pipes (inner diameter: 68 mm; outer diameter: 76 mm) were vertically connected to reach one of four target heights (2, 3, 5, or 9 m), either inside the laboratory or along the outer wall of the laboratory building (at each joint, the pipe wall formed a double-layered structure; see Figure 1). The apparatus was filled with dechlorinated tap water stored for one day prior to use.

**Figure 1.**
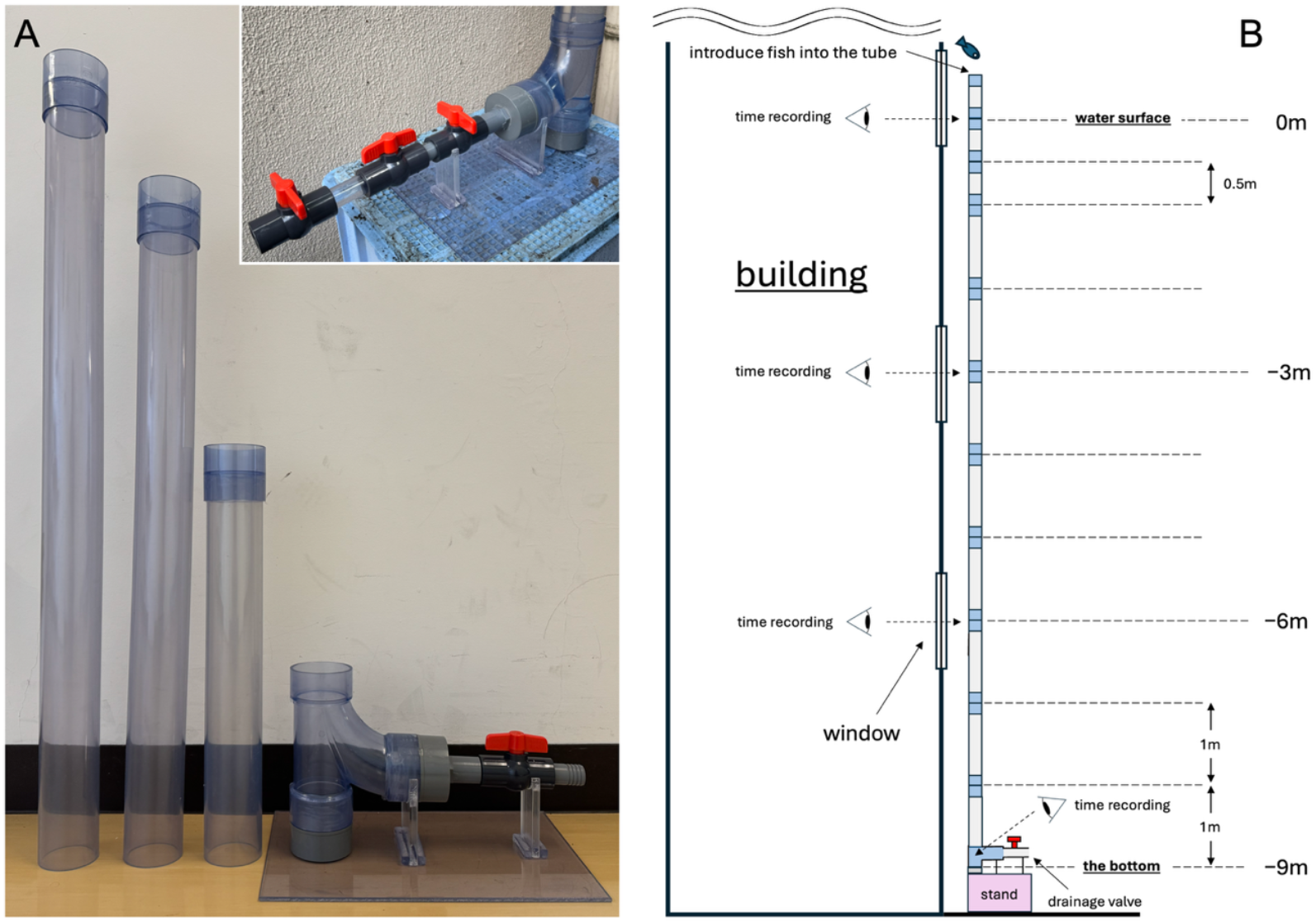
Experimental cylindrical tank used in this study. A. Components of the PVC tank (before assembly): three types of transparent PVC pipes and the bottom structure. A drainage valve is attached at the base. The pipes on the left and right are 1 m and 50 cm in length, respectively; the central piece, when connected to the bottom structure, reaches exactly 1 m in height. Inset: The adapter attached to the base of the apparatus, which was used to introduce fish from the bottom of the tank (see main text). B. Schematic of the tank after assembly. Fish were introduced from the top, and the times at which they passed the 3 m and 6 m marks were recorded by visual observation. For the 9 m mark, the recorded time corresponds to the moment the fish entered the bottom structure, not when it made contact with the tank floor.

For indoor setups, the water temperature was approximately 25°C, and illumination was provided by ceiling-mounted fluorescent lights. For outdoor setups, experiments were conducted under natural sunlight, and the initial water temperature was ∼25°C; the water temperature at the end of the experiment was not recorded but might have increased because the experiments were conducted on sunny summer days.

A single adult fish was introduced into the water column, and the time required to reach a specified target depth for the first time was recorded. Depth-specific observations were made through building windows located at 3, 6, and 9 m above ground level (corresponding to the 1.5th, 2.5th, and 3.5th floors, respectively), with the 9 m depth observed from the ground. Once the fish reached the maximum depth, or was judged unlikely to do so, the next fish was introduced and the procedure repeated. If a previously observed individual resurfaced during the experiment and individual identification remained possible, the trial was continued. If multiple individuals overlapped and identification became unreliable, the experiment was terminated; the water was drained, the fish were retrieved, and the column was refilled before resuming observations with untested individuals.

### Ethics statement

All experimental procedures involving fish were conducted in accordance with the institutional guidelines for animal care and use and were approved by the Animal Experiment Committee of Japan Women’s University.

## Results

### Preliminary experiment using medaka

A preliminary trial was conducted at a depth of 2 m inside the laboratory, based on the initial assumption that medaka would not voluntarily dive to such depths. Three male medaka of mixed genetic background (n = 3) were tested individually. All exhibited hesitation or reversed direction at the pipe junctions during descent—a behavior observed in subsequent experiments—but ultimately reached the bottom of the tank without coercion (unpublished observation).

The experiment was then conducted outdoors at a depth of 3 m using ten wild medaka from Nakaikemi (n = 10), introduced individually. All reached the bottom within 5 min and remained at maximum depth for the entire observation period (up to ∼30 min) without resurfacing (unpublished observation).

After draining the tank, the pipes were extended to a depth of 5 m. In a repeated trial with the same Nakaikemi population (n = 10, which could partially overlap with the previous trial), all individuals reached the bottom within 7 min. One fish left the bottom during the ∼40 min observation and repeatedly ascended and descended, but no other individuals displayed following, avoidance, or aggression toward this behavior (unpublished observation).

These preliminary results suggested that medaka are capable of voluntarily diving to depths of at least 5 m, prompting more systematic experiments to examine this behavior under controlled conditions.

### Main experiment using medaka

The results of the 9-m outdoor experiment (Figure 1B) are shown in Figure 2A. All d-rR strain individuals (n = 10) voluntarily reached the bottom. The times to reach depths of 3, 6, and 9 m were 1 min 3 s ± 12 s, 2 min 7 s ± 26 s, and 2 min 41 s ± 31 s (mean ± SE), respectively. No significant sex differences were detected at any depth (n = 5 each; p ≥ 0.297, Student’s two-tailed t-test).

**Figure 2.**
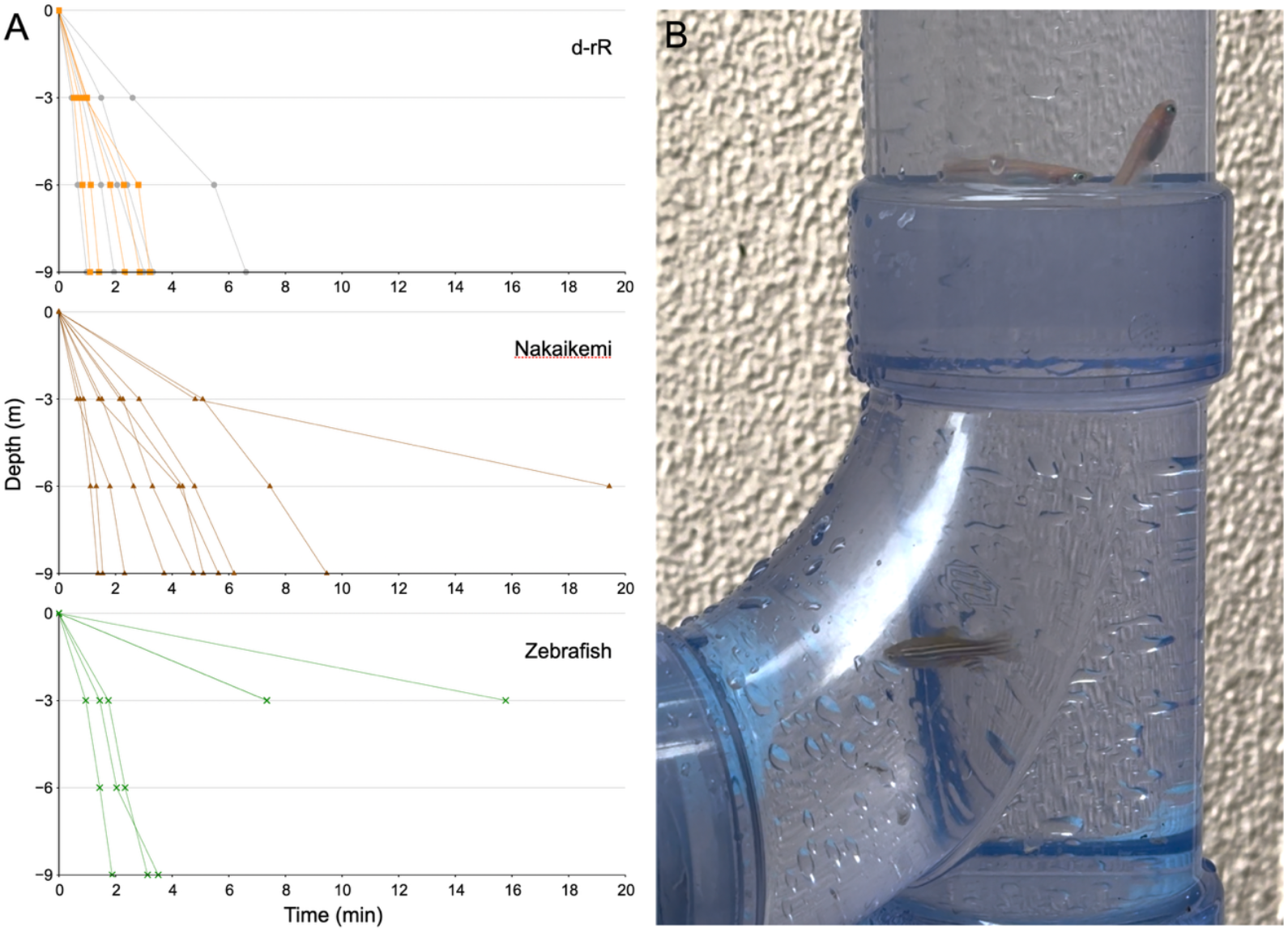
Experimental results at a depth of 9 meters. A. Time required for individual fish to reach depths of 3, 6, and 9 meters. Top: d-rR; middle: Nakaikemi; bottom: zebrafish. Data from all individuals are shown. All 10 d-rR individuals reached 9 meters. Females and males are distinguished by gray and orange lines, respectively. For Nakaikemi, one individual did not reach 9 meters, resulting in a truncated line. For zebrafish, a maximum of six individuals reached 3 meters, and three reached 9 meters (see main text). Two of the six individuals that reached 3 meters did so simultaneously (at 7 min 21 sec), resulting in overlapping lines. B. Visual snapshot of the 9-meter tank. One zebrafish is visible near the bottom, while two medaka are seen near the top.

For the Nakaikemi group (n = 10), nine individuals voluntarily reached the bottom. The times to reach depths of 3, 6, and 9 m were 1 min 57 s ± 28 s, 3 min 27 s ± 40 s, and 4 min 27 s ± 51 s (mean ± SE), respectively. These times tended to be longer than those of the d-rR strain, but the differences were not statistically significant at any depth (p > 0.079). The remaining individual repeatedly ascended and descended, reaching 6 m after 19 min 26 s but failing to reach the bottom during the >40-min observation period.

For both d-rR and Nakaikemi medaka, the descent time from −6 m to −9 m (35 s ± 6 s and 1 min 0 s ± 12 s, respectively) was clearly shorter than from 0 m to −3 m (1 min 3 s ± 12 s and 1 min 57 s ± 28 s, respectively) and from −3 m to −6 m (1 min 3 s ± 15 s and 1 min 30 s ± 17 s, respectively). This trend may reflect reduced buoyancy due to swim bladder compression under increased hydrostatic pressure. However, the differences were not statistically significant (p = 0.163 for both groups, one-way ANOVA).

### Zebrafish diving behavior

For zebrafish (n = 10), four individuals were introduced first. The first and third reached the bottom at 3 min 8 sec and 3 min 31 sec, respectively, but resurfaced during the experiment. The second failed to reach 6 meters after 24 minutes, and the fourth did not reach 3 meters within 18 minutes. Once the fish resurfaced, it became impossible to distinguish them from each other, so their subsequent behavior could not be tracked individually.

An attempt was made to retrieve all four fish from the surface using a net, but only one (identity unknown) was caught. Without draining the column, the remaining six zebrafish were introduced while three unidentified fish from the first group were still in the tank.

Among the newly introduced fish, one individual reached the bottom in 1 min 53 sec, and two others reached 3 meters at 7 min 21 sec. However, it could not be determined whether these were from the first or second batch. Figure 2B presents the data assuming they were from the second batch. Under this assumption, the maximum numbers of fish reaching 3, 6, and 9 meters were 6, 3, and 3, respectively; the corresponding minimum numbers (if some were from the first batch) were 3, 2, and 2. In either case, the results contrasted sharply with medaka, in which nearly all individuals reached 9 meters. The six fish that did not dive remained near the surface until the experiment was terminated after an additional 12 minutes.

### Ascending behavior

Finally, we conducted experiments in which fish were introduced from the bottom of the tank using an adapter equipped with stopcocks on both sides (Figure 1A). The adapter was filled with water, a fish was confined inside, and the adapter was connected to the base of the device (drainage section; Figure 1A). After opening the stopcocks on both sides of the connection and allowing the fish to move into the base, observations were initiated. Covering the transparent part of the adapter with aluminum foil appeared to facilitate the movement of fish toward the base (the brighter region).

The water depth was increased to 10 m by adding an additional 1-m pipe segment, and zebrafish were tested first (n = 10; individuals distinct from those used above). The first fish ascended to the water surface after 13 min 47 sec and was retrieved from the top using a net. The second fish similarly reached the surface after 18 min 32 sec and was also removed. The third fish failed to reach even 8 m after 14 minutes; therefore, a fourth fish was introduced. These two fish then tended to move together and repeatedly ascended and descended. However, even 19 minutes after the introduction of the fourth fish (33 minutes after the introduction of the third), the maximum ascent reached was approximately 7 m. An attempt to recover the fish by draining the tank when they approached the bottom resulted in the retrieval of only one individual.

After replenishing the drained water, an additional six zebrafish were introduced from the bottom. The remaining seven fish tended to act collectively, but again showed only repeated vertical movements (Movie 2). Four individuals ascended to a depth of 4 m after 12 min 42 sec, but none reached the water surface, and observations were terminated 28 minutes after introduction.

After draining the tank and recovering the fish, the same experimental procedure was applied to medaka (Nakaikemi; n = 10). Based on previous observations suggesting that medaka are less prone to ascend than zebrafish, individuals were introduced one by one at intervals of approximately 10 minutes. After three individuals had been introduced, all remained near the bottom, although one fish ascended to a depth of 7.5 m at 32 minutes after the start of the experiment. The fourth fish failed to exit the adapter; therefore, the adapter was removed, and the remaining six individuals (total n = 7) were introduced simultaneously at 40 minutes. One individual ascended to 7 m after 57 minutes and subsequently to 4 m after 61 minutes, whereas the remaining nine individuals stayed near the base throughout the observation period (as shown in Movie 1), with a maximum ascent depth of 7 m. The individual that reached 4 m did not continue ascending and instead began to descend; observations were therefore terminated 63 minutes after the start of the experiment.

## Discussion

It is fair to acknowledge the criticism that the behavior observed in this study may have occurred only under the highly artificial conditions of a narrow-diameter tank. It is also likely that environmental conditions such as water temperature, light intensity, turbidity, and current would influence their behavior. Nevertheless, it is important to emphasize that, with the exception of a single individual, all medaka reached the tank bottom at all tested depths, despite the fact that the setup allowed for the alternative behavior of remaining near the surface, as observed in many zebrafish. What can be stated with certainty based on this study is that medaka possess the intrinsic ability to voluntarily dive to depths of at least 9 meters and remain there for extended periods.

It is well known that when fish are introduced into an unfamiliar environment, they tend to immediately descend to the bottom and remain relatively immobile [Johnson et al., 2023]. The 9-meter-deep cylindrical tank used in this study was undoubtedly a novel environment for the medaka, and they reached the bottom quickly and remained there. However, in medaka, anxiety-induced bottom-dwelling typically lasts no more than one minute. After that initial minute, medaka begin to explore their surroundings, and once they have become familiar with the environment, they tend to relax and exhibit reduced movement [Matsunaga & Watanabe, 2010]. In this study, observations after reaching the deepest point showed that the fish were swimming around somewhat actively, rather than freezing in a rigid, anxious state (see Video 1). Therefore, although the initial descent may have been triggered by anxiety due to being placed in a novel environment, the continued stay at the bottom likely had a different underlying cause.

As one possible hypothesis, we propose that medaka prefer environments where the bottom is visually accessible. Medaka have generally been regarded—as we ourselves also believed—as surface-dwelling fish favoring the uppermost layer of the water column. However, under all tested conditions, the consistent tendency of the fish to remain near the bottom—far from the surface—suggests a preference for the benthic environment. Perhaps medaka are simultaneously attracted to both the water surface and the bottom, and this dual preference may explain their common occurrence in shallow waters where both are within reach. Medaka may be less frequently distributed in deeper habitats characterized by surface-only zones (e.g., the upper layer of deep water), bottom-only zones, or midwater zones where neither surface nor bottom is visible. Nevertheless, our original question—”How deep should we extend our search when surveying wild medaka populations in wetlands with depths up to 4 meters?”—has led us to the unequivocal conclusion that the search depth should not be restricted.

For zebrafish, the relatively small number of individuals that reached the bottom (Figure 2A) suggests that their preference for the bottom is weaker than that of medaka. Consistent with this interpretation, in the bottom-introduction experiments, two individuals ascended all the way to the water surface, while the remaining eight repeatedly alternated between ascending and descending movements (Movie 2). The precise reason why most individuals ultimately failed to reach the surface remains unclear. However, the observation that zebrafish frequently reversed direction and began descending at depths of approximately 4–7 m (Movie 2) raises the possibility that they sensed some form of physiological disturbance associated with changes in hydrostatic pressure, potentially involving swim bladder expansion. It is conceivable that, had the experiments been conducted at shallower depths (e.g., 4–7 m), zebrafish might have reached the surface and remained there for extended periods.

The device we developed (Figure 1) can be extended to greater depths by connecting additional pipe segments, provided that structural stability is ensured. Although currently limited to small fish species or juvenile and larval stages, this apparatus holds promise for future studies employing a range of species—such as surface-dwelling hatchetfish or benthic gobies— offering novel insights into fish behavior, physiology, and ecology from previously unexplored perspectives.

## Figure/Movie Legends

**Movie 1. Medaka schooling near the bottom at a depth of 9 m**.

Each individual was introduced into the tank separately, but since they remained near the bottom after reaching the deepest point, they eventually gathered closely together. Although their movements appear somewhat restless, all fish are actively swimming and show no signs of immobility or trembling associated with freezing behavior.

**Movie 2. Zebrafish repeatedly ascending and descending at depths of 6–10 m**.

Seven zebrafish were introduced from the bottom of the tank. The fish actively repeated ascending and descending movements without showing any apparent signs of damage caused by increased hydrostatic pressure; however, none of the individuals reached the water surface.

## Acknowledgements

We are deeply grateful to Mr. Toshiyuki Saji of the National Institute for Physiological Sciences for his invaluable contribution in designing and constructing the cylindrical water column used in this study. We also thank Dr. Takako Yasuda of Japan Women’s University for kindly providing the d-rR strain. Our gratitude also goes to the undergraduate students of Japan Women’s University—E. Kondo, A. Kumagai, K. Odamaki, and M. Takemura—for their assistance during the 3-, 5-, 9-, and 10-meter experiments.

## Data Availability

The datasets used and/or analyzed during the current study available from the corresponding author (S.F.) on reasonable request.

